# The Claustrum Coordinates Cortical Slow-Wave Activity

**DOI:** 10.1101/286773

**Authors:** Kimiya Narikiyo, Rumiko Mizuguchi, Ayako Ajima, Sachiko Mitsui, Momoko Shiozaki, Hiroki Hamanaka, Joshua P. Johansen, Kensaku Mori, Yoshihiro Yoshihara

## Abstract

During sleep and awake rest, the neocortex generates large-scale slow-wave activity. Here we report that the claustrum, a poorly understood subcortical neural structure, coordinates neocortical slow-wave generation. We established a transgenic mouse line allowing genetic and electrophysiological interrogation of a subpopulation of claustral glutamatergic neurons. These claustral excitatory neurons received inputs from glutamatergic neurons in a large neocortical network. Optogenetic activation of claustral neurons *in vitro* induced excitatory post-synaptic responses in most neocortical neurons, but elicited action potentials primarily in inhibitory interneurons. Optogenetic activation of claustral neurons *in vivo* induced a Down-state featuring a prolonged silencing of neural acticity in all layers of many cortical areas, followed by a globally synchronized Down-to-Up state transition. These results demonstrate a crucial role of the claustrum in synchronizing inhibitory interneurons across the neocortex for spatiotemporal coordination of brain state. Thus, the claustrum is a major subcortical hub for the synchronization of neocortical slow-wave activity.

---

During sleep and awake rest states, multiple cortical areas synchronously generate slow-wave (SW) activity critical for brain synaptic homeostasis and the replay and consolidation of memory traces acquired during the preceding waking epoch (*1-4*). The generation of SW activity involves a silent Down-state and active Up-state (*5*). However, the neural structures and mechanisms that control SW activity across large-scale cortical areas remain unknown.

The claustrum is a thin, sheet-like neural structure located between the insular cortex and the striatum, with reciprocal connections to and from nearly all neocortical areas (*6*), that has been hypothesized to play crucial roles in higher brain functions such as consciousness, multisensory integration, salience detection, and attentional load allocation (*7-9*). However, there have been only a few experimental studies of claustrum function (*10,11*), and its precise roles in brain physiology have not been elucidated. In this study, we developed claustrum-specific transgenic tools in the mouse, and show that a population of claustrum excitatory neurons regulates SW activity over widespread cortical areas in a state-dependent manner by the synchronized silencing of cortical neurons.

## Results

### Claustrum-specific Cre-expressing Transgenic Mouse Line

In the course of our olfactory system research generating transgenic mouse lines with the mitral/tufted cell-specific *Tbx21* gene promoter (*12*), we serendipitously found an interesting line in which Cre recombinase was ectopically and specifically expressed in the claustrum of adult brain (Fig. 1a). Cre-positive neurons were distributed along the entire antero-posterior and dorso-ventral axes of the claustrum, but not in other brain regions. Higher magnification revealed that Cre-positive neurons were mostly confined to the claustrum, while only a small number of Cre-positive neurons were detected in neighboring regions such as the dorsal endopiriform nucleus and the deepest part of the insular and somatosensory cortex (Fig. 1b). The claustrum-specific Cre-expressing transgenic mouse line was designated Cla-Cre and it enabled genetic visualization and manipulation of those neurons.

**Fig. 1.**
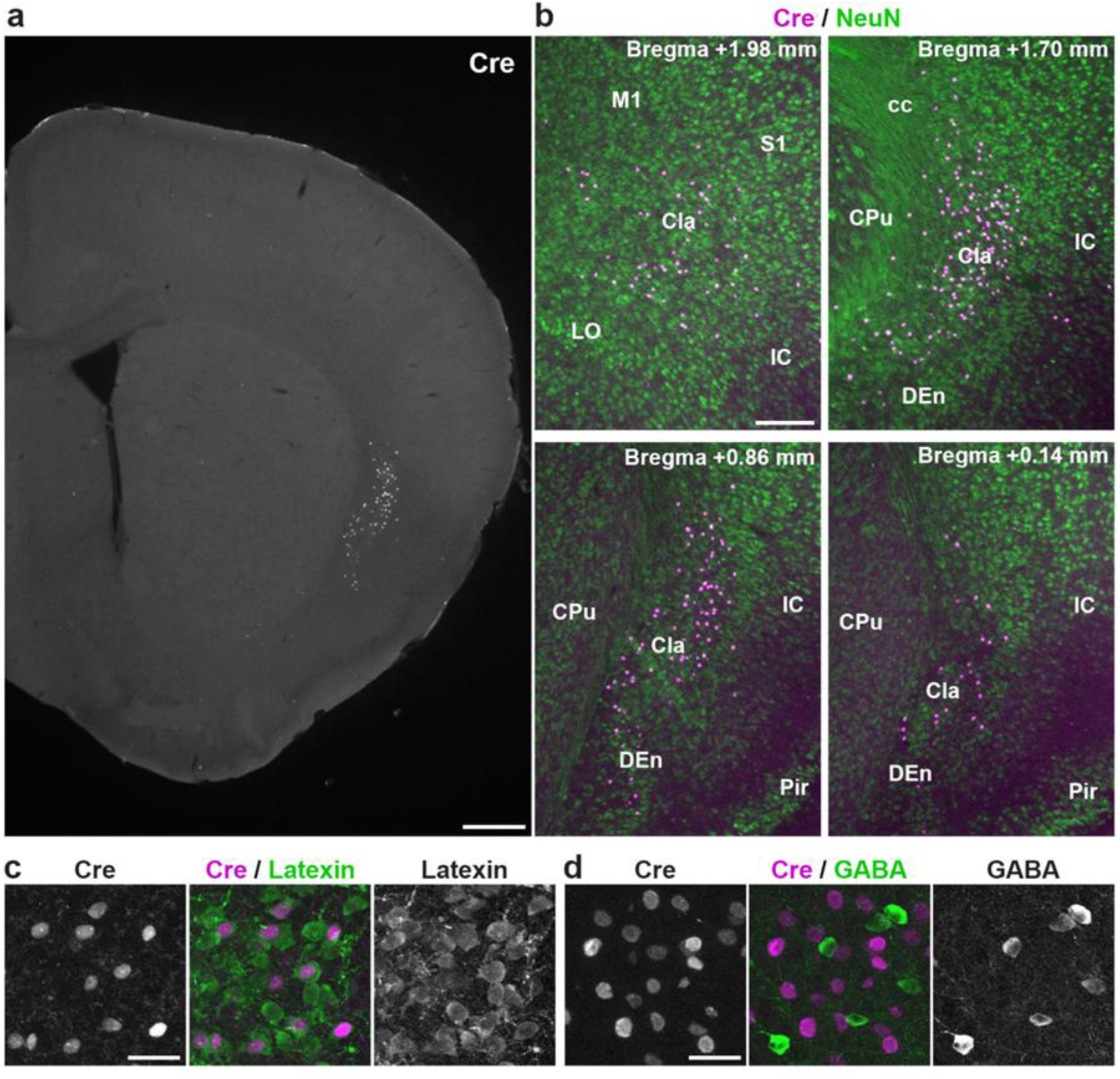
Characterization of Cre-expressing Neurons in Cla-Cre Transgenic Mice. **a**, Expression of Cre recombinase in the Cla-Cre mouse. A coronal section of Cla-Cre mouse brain was immmunostained with an anti-Cre antibody. The Cre-expressing neurons are present specifically in the claustrum. Scale bar, 500 µm. **b**, Higher magnification of the claustrum (Cla) and surrounding regions. Coronal sections at different levels along the antero-posterior axis of the Cla-Cre mouse brain were double-stained with anti-Cre (magenta) and anti-NeuN (green) antibodies. The Cre-expressing neurons are observed in a subset of claustral neurons. cc, corpus callosum; CPu, caudate putamen; DEn, dorsal endopiriform nucleus; IC, insular cortex; LO, lateral orbital cortex; M1, primary motor cortex; Pir, piriform cortex; S1, primary somatosensory cortex. Scale bar, 200 µm. **c**, Double immunofluorescence staining of the claustrum in the Cla-Cre mouse with anti-Cre (magenta) and anti-Latexin (green) antibodies. Cre-expressing neurons are positive for Latexin, a marker of glutamatergic neurons in the claustrum, indicating that they are glutamatergic excitatory neurons. Scale bar, 30 µm. **d**, Double immunofluorescence staining of the claustrum in the Cla-Cre mouse with anti-Cre (magenta) and anti-GABA (green) antibodies. All the Cre-positive neurons are negative for GABA, indicating that they are not GABAergic inhibitory neurons. Scale bar, 30 µm.

The claustrum contains both glutamatergic excitatory neurons and GABAergic inhibitory neurons (*13*). To determine which type of claustral neurons expressed Cre recombinase, we immunohistochemically characterized the Cre-positive neurons with anti-Latexin and anti-GABA antibodies. Latexin is a marker of glutamatergic neurons in the claustrum (*14*). Cre was expressed in about one-third (34.6 ± 2.3 %, n = 3 mice) of Latexin-positive glutamatergic neurons (Fig. 1c). In contrast, Cre-positive neurons did not show any GABA signals (Fig. 1d). Thus, Cre labels a major subset of glutamatergic neurons in the claustrum.

### Claustral Neuronal Inputs Comprise a Large Cortical Excitatory Network

First, we genetically visualized neuronal inputs to the claustrum in Cla-Cre mice. To label presynaptic neurons projecting axons to Cre-positive claustral neurons, we used a trans-synaptic retrograde tracing method combining a modified rabies virus (RVΔG) with an adeno-associated virus (AAV)-mediated Cre-dependent expression of the EnvA receptor (TVA) and rabies glycoprotein (RG) (*15*) (Fig. 2a,d,e and Supplementary Fig. 1). For quantification, we counted the number of GFP-positive presynaptic neurons in individual brain areas (Fig. 2c, left; n = 4 mice). Claustral neurons received dense inputs from the agranular insular cortex, a cortical area that overlies the claustrum. In addition, the claustrum was heavily innervated by neurons in many higher-order cortical areas, such as the orbital, prelimbic, cingulate, secondary motor, granular insular, and entorhinal cortices. Among sensory cortical areas, the somatosensory and olfactory cortices showed substantial innervation to the claustrum, whereas relatively sparse inputs were observed from the visual and auditory areas. Many presynaptic neurons were also present in the prefrontal cortex of the contralateral hemisphere (Fig. 2a). The claustrum-targeting neurons in the cortex were abundantly located in layer 5, and many showed pyramidal morphology with apical and basal dendrites (Fig. 2d and Supplementary Fig. 1). Several subcortical structures also provided synaptic inputs to the claustrum, including the basolateral amygdala, the thalamic mediodorsal and parafascicular nuclei, the substantia innominata, and the raphe nuclei (Fig. 2, a, d, e, f and Supplementary Fig. 1). The input neurons to the claustrum were glutamatergic, except for cholinergic neurons in the substantia innominata, serotonergic neurons in the raphe nuclei, and some GABAergic neurons within the claustrum (Fig. 2d,e). No innervation was observed from dopaminergic neurons in the ventral tegmental area or noradrenergic neurons in the locus coeruleus (data not shown).

**Fig. 2.**
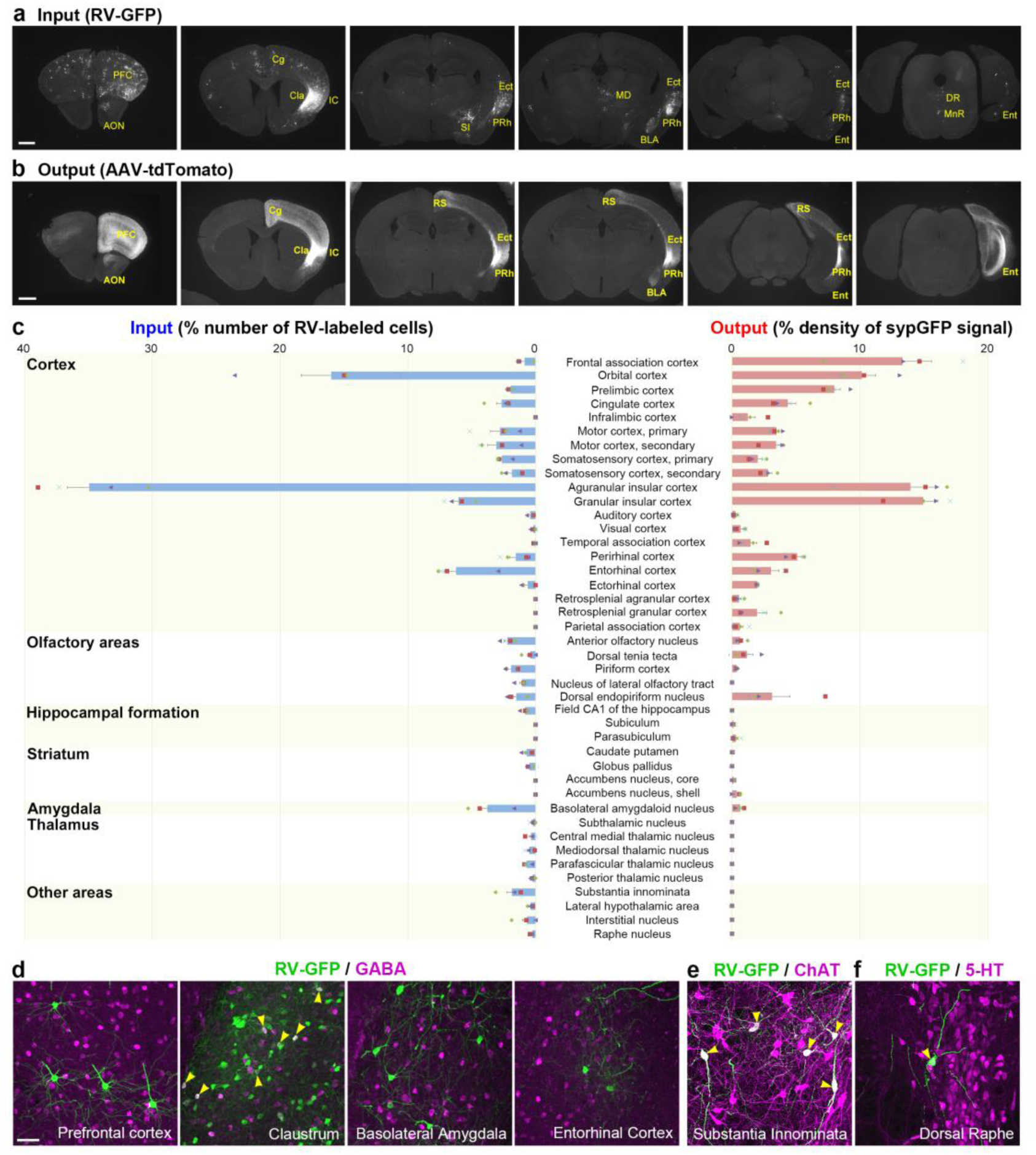
Genetic Neural Circuit Mapping of Claustrum Input-Output Patterning. **a**, Presynaptic input neurons to Cre-expressing claustral neurons were visualized with a modified rabies virus-mediated mono-synaptic retrograde tracing method. GFP-positive presynaptic neurons are distributed in various neocortical areas as well as the basolateral amygdala (BLA), substantia innominata (SI), thalamic mediodorsal nucleus (MD), dorsal raphe nucleus (DR) and median raphe nucleus (MnR). AON, anterior olfactory nucleus; Cg, cingulate cortex; Cla, claustrum; Ect, ectorhinal cortex; Ent, entorhinal cortex; IC, insular cortex; PFC, prefrontal cortex; PRh, perirhinal cortex. Scale bar, 1 mm. **b**, Axonal trajectories of Cre-expressing claustral neurons were visualized with Cre-dependent tdTomato-expressing AAV. Claustral axons are observed in various neocortical areas as well as in the BLA. RS, retrosplenial cortex. Scale bar, 1 mm. **c**, Quantification of the input neurons and the output areas of the Cre-expressing claustral neurons. (Left) Numbers of GFP-positive presynaptic neurons in individual brain regions were counted on sections labeled with GFP-expressing modified rabies virus. (Right) Cre-dependent synaptophysin-GFP-expressing AAV was injected into the claustrum of Cla-Cre mice and densities of synaptophysin-GFP-positive puncta were measured in each brain area. Error bars are s.e.m. **d-f**, Double immunofluorescence staining of GFP-positive input neurons (green) with neuronal type markers. Magenta signals represent GABA (**d**), choline acetyltransferase (**e**) and serotonin (**f**). GABAergic presynaptic neurons are observed only within the claustrum (**d**). The claustrum receives neuromodulatory inputs from cholinergic neurons in the substantia innominata (**e**) and serotonergic neurons in the raphe nucleus (**f**). Yellow arrowheads denote double-positive neurons. Scale bar, 50 µm.

### Claustral Neuronal Outputs Define a Widespread Cortical Network

To investigate the target brain regions innervated by Cre-positive claustral neurons, we examined their axonal projections and synaptic vesicle distributions by AAV-mediated Cre-dependent expression of tdTomato (Fig. 2b and Supplementary Fig. 2) and synaptophysin-GFP (Fig. 2c and Supplementary Fig. 3), respectively. tdTomato-expressing claustral axons were mostly confined to, but broadly and diffusely distributed within the cerebral cortex. Except for the claustrum itself, only a few subcortical structures received claustral projections, such as the dorsal endopiriform nucleus and the basolateral amygdala. To compare the degree of claustral innervation of these brain regions, we quantified the densities of synaptophysin-GFP signals and plotted the output proportions (Fig. 2c, right; n = 4 mice). The highest degree of claustral innervation was observed in the insular, frontal association, orbital, and prelimbic cortices, followed by other higher-order cortical areas such as the cingulate, secondary motor, perirhinal, entorhinal, ectorhinal, and retrosplenial cortices. In each area, the synaptophysin-GFP signals were distributed in all the layers (Supplementary Fig. 3). The somatosensory cortex and several olfactory cortical areas also received substantial axonal projections from the claustral neurons, while claustral projections to the visual and auditory cortices were more sparse.

**Fig. 3.**
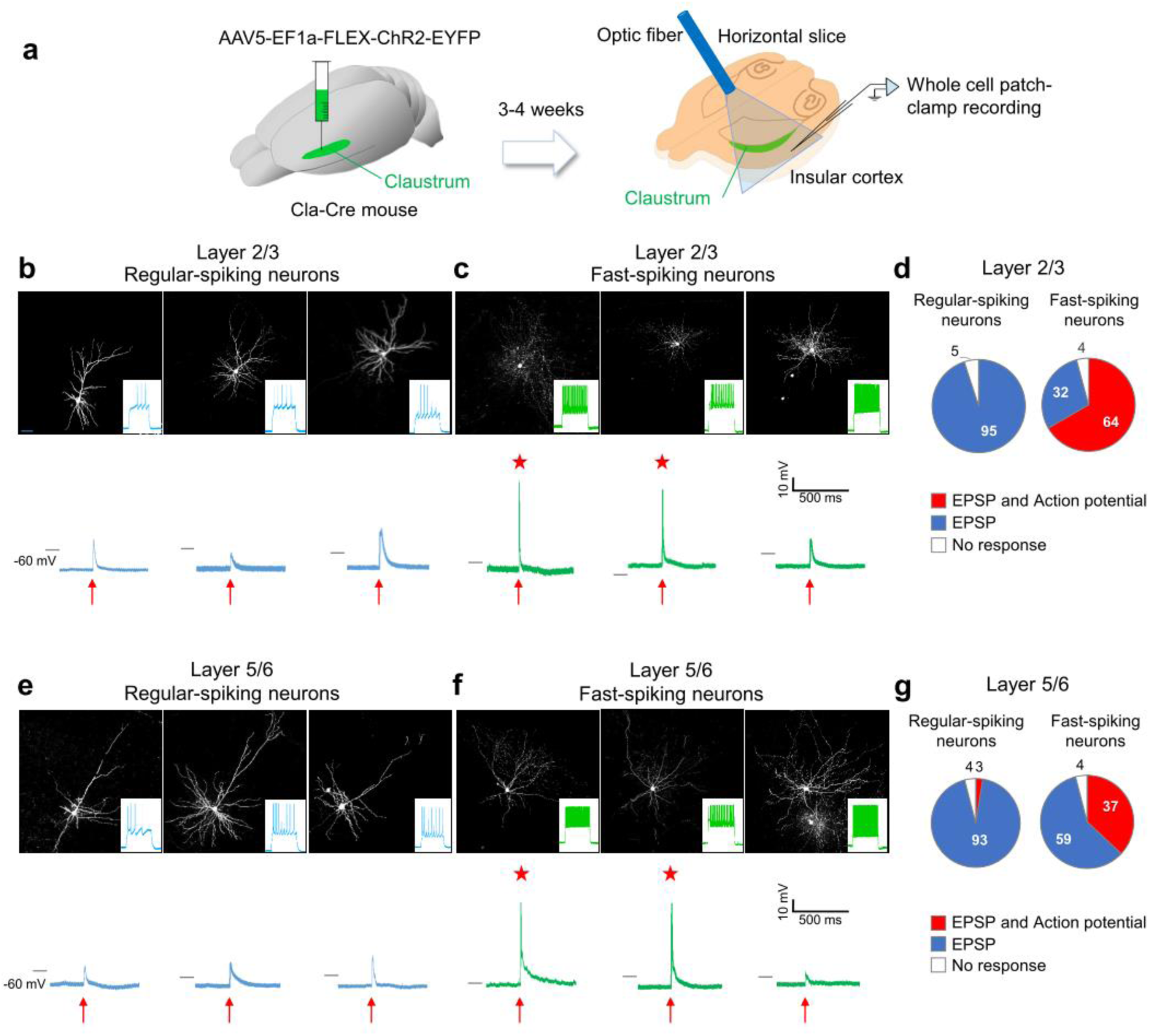
Claustral Activation Drives a Cortical Fast-Spiking Interneuron Network. **a**, Schematic of *in vitro* whole cell patch-clamp recording from insular cortical neurons with claustral optogenetic stimulation **b**, Current-clamp recordings of layer 2/3 regular-spiking neurons. The morphology of recorded neurons filled with biocytin (top), firing patterns induced by inward current pulse injection (inset), and responses to optogenetic stimulation of claustral axons (bottom). **c**, Recordings from the layer 2/3 fast-spiking neurons. **d**, Percentage of layer 2/3 regular-spiking neurons (left, n = 38) and fast-spiking neurons (right, n = 28) that showed EPSP with action potential (red), EPSP without action potential (blue), and no response (white) upon optogenetic claustral stimulation. **e**, Recordings from layer 5/6 regular-spiking neurons (n = 76). **f**, Recordings from layer 5/6 fast-spiking neurons (n = 27). **g**, Percentage of layer 5/6 regular-spiking neurons (left, n = 76) and fast-spiking neurons (right, n = 27) that showed an EPSP with action potential (red), EPSP without action potential (blue), and no response (white) upon the optogenetic claustral stimulation. Black bar on the left of each voltage trace, −60 mV; red arrow below each trace, light stimulation (5 ms); red asterisk, action potential.

### In Vitro Optogenetic Activation of Claustal Neurons Drives Cortical Interneurons

Based on the widespread axonal projections from Cre-positive claustral neurons to the cortex, we asked what types of neuronal responses were evoked in cortical neurons upon optogenetic stimulation of the claustrum. Cre-dependent channelrhodopsin2 (ChR2)-expressing AAV was injected into the claustrum of Cla-Cre mice. After three to four weeks, we prepared horizontal brain slices containing both the claustrum and the neighboring agranular insular cortex and performed whole-cell patch-clamp recordings from insular neurons in combination with photo-stimulation of claustral neurons (Fig. 3a). The recorded insular neurons were categorized into four classes according to their location in cortical layers 2/3 or 5/6 and regular-spiking or fast-spiking firing patterns after a depolarizing current pulse injection. Most regular-spiking neurons harbored a pyramidal-shaped soma with apical and basal spiny dendrites and produced wide-spike waveforms, consistent with their identity as excitatory pyramidal neurons (Fig. 3b,e) (*16*). In contrast, the fast-spiking neurons showed narrow-spike waveforms and a complex dendritic arborization, indicating that they represent a subset of inhibitory interneurons (Fig. 3c,f) (*17,18*).

Photo-stimulation of ChR2-expressing claustral neurons evoked excitatory post-synaptic potentials (EPSPs) in almost all recorded neurons, including both the regular- and fast-spiking neurons in all of the layers (Fig. 3d,g). However, action potential responses were observed frequently in fast-spiking neurons (64% in layer 2/3; 37% in layer 5/6), but rarely in regular-spiking neurons (0% in layer 2/3; 2.6% in layer 5/6). These findings suggest that Cre-positive claustral neurons induce EPSPs in all types of neurons but drive spike responses predominantly in a subset of inhibitory interneurons in the insular cortex. Similar results were obtained in neuronal recordings in the frontal cortex (Supplementary Fig. 4).

**Fig. 4.**
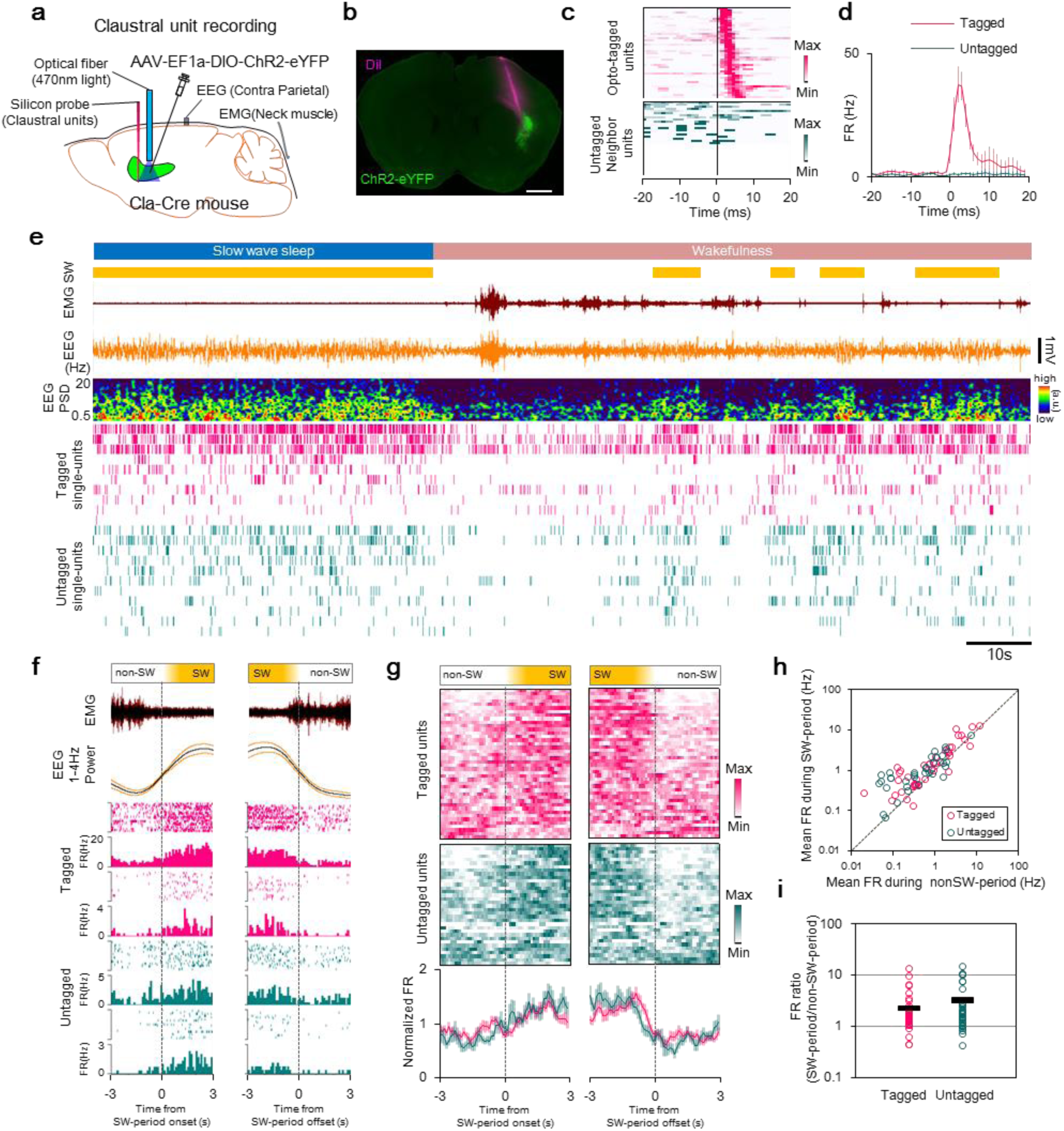
Claustral Neurons are Active during Neocortical Slow-Wave Activity. **a**, Schematic of claustrum unit recording in head-fixed mice. **b**, An example of the histological identification of claustrum recording sites. Claustral neurons express ChR2-eYFP (green). DiI (magenta) shows the recording electrode track. Scale bar, 1 mm. **c**, Photo-stimulation-induced spike responses of opto-tagged units (magenta, n = 43) and neighboring untagged units (blue, n = 34). Each line indicates color-coded peri-stimulus time histogram (PSTH, 1-ms bins, smoothed by 3 bins moving average) of spike activity of singly isolated unit. Opto-tagged units show spike activity with latencies within 5 ms after the photo-stimulation onset. **d**, Group averages of the PSTHs of the photo-stimulation-induced spike responses. Magenta line, opto-tagged unit group; Blue line, untagged unit group. FR, firing rate. Error bars are s.e.m. **e**, An example of claustral single-unit activities and simultaneously recorded EMG and EEG during the transition from SW sleep to wakefulness. Yellow bar indicates the period when EEG shows SW activity (1-4 Hz) (SW-period). PSD, power spectral density. **f**, Examples of averaged EMG, SW activity (EEG 1-4 Hz power) and peri-event time histograms (PETHs, 0.1-s bins) of representative claustral units in reference to the onset time of SW-period (left column) and the offset time of SW-period (right column). **g**, Color-coded PETHs of spikes of claustral units (magenta, opto-tagged units; blue, untagged units) aligned in reference to the onset time of SW-period (left) and the offset time of SW-period (right). The bottom histograms show averages of the PETHs. Each PETH was calculated using 23 to 30 events of onsets or offsets of SW-period, and smoothed by a moving average of 3 bins. For averaging analysis, each FR was divided by its mean FR to normalize values. Error bars, s.e.m. Spike activity around the onset of SW-period (left) did not differ significantly between groups (interaction, F(59,4366) = 1.12, P = 0.24, two-way repeated-measures ANOVA). In spike activity around the offset of SW-period (right), two-way repeated-measures ANOVA detected interaction (F(59,4366) = 1.49, P = 0.0088), but *post hoc* test did not detect significant difference between groups at any time bin. **h**, Scatter plot of mean FR (Hz) of each unit in **g** during SW-period vs. non-SW-period. The mean FR of non-SW-period was calculated by averaging FR during −3 to −1 s from the onset of SW-period and 1 to 3 s after the offset of SW-period. For SW-period, data of 1 to 3 s after the onset of SW-period and −1 to −3 s from the offset of SW-period were used. **i**, FR ratio of each unit between SW-period and non-SW-period (SW-period/non-SW-period). FR ratios were calculated based on **h**. The mean FR ratios were significantly higher than 1 in both opto-tagged (mean ± s.e.m., 2.3 ± 0.4, *t*(41) = 3.4, P = 0.0014, two-sided one sample *t* test) and untagged (mean ± s.e.m., 3.3 ± 0.6, *t*(33) = 3.7, P = 0.00078, two-sided one sample *t* test) groups. There are no significant defference between groups (*t*(56.8) = −1.3, P = 0.19, two-sided Welch’s two sample *t* test).

### Slow-Wave-associated Firing of Claustral Neurons in Sleep and Resting Wake

We examined whether claustral neuron activity is brain-state dependent by measuring their firing patterns during sleep and wake states. We conducted *in vivo* electrophysiological recordings of claustral neuron spike activity in head-fixed, unanesthetized Cla-Cre mice using linear silicon multi-channel microelectrode arrays (Fig. 4a). Parietal cortex surface electroencephalography (EEG) and neck muscle electromyography (EMG) were simultaneously recorded to monitor behavioral state. Spike activities of Cre-positive claustral neurons were identified through opto-tagging following AAV-mediated ChR2 expression (Fig. 4b-d). Opto-tagged claustral units (Tagged, n = 42 units from 5 mice) and their neighboring untagged units, which are presumably spike activities from Cre-negative neurons in the claustrum (Untagged, n = 34 units from 5 mice), were analyzed separately (Fig. 4e). The majority of claustral neurons in both groups increased their firing rate during the period when the neocortical EEG showed SW (1–4 Hz) activity (SW-period) compared with the period when the EEG did not show SW activity (non-SW-period). The SW-periods occured during SW sleep (continuous low EMG and SW-dominant periods) and quiet wakefulness (transient low EMG periods between high EMG periods), but rarely during active wakefulness (high EMG periods) (Fig. 4e). Importantly, claustral neuronal activity depended on the presence of SW activity. Peri-event time histograms of claustral unit activities in relation to the onset or offset of SW-period (Fig. 4f,g) confirmed that a majority of claustral neurons were more active during the SW-period than non-SW-period. Comparison of mean firing rates between SW-periods and non-SW-periods of each neuron revealed that most claustral neurons showed higher firing rates during SW-periods (SW-period firing rate / non-SW-period firing rate > 1; Tagged: 38 / 42 units [90%]; Untagged: 28 / 34 units [82%]) (Fig. 4h,i). The increased activity during SW-periods together with the widespread projections of claustral neurons to the cortex suggest the possibility that claustral neuron activity is involved in the regulation of cortical SW activity.

### In Vivo Optogenetic Activation of Claustral Neurons Triggers a Cortical Down State

To clarify the causal relationship between claustral excitatory neuron activity and neocortical SW generation, we investigated the effect of optogenetic claustral activation on neocortical neuron firing state *in vivo*. Both LFP and unit activity were recorded in frontal cortex (FC) regions including frontal association cortex, orbitofrontal cortex, and secondary motor cortex, which receive dense inputs from claustral neurons (Fig. 2b,c), while ChR2-expressing claustral neurons were photo-stimulated (Fig. 5a,b). During SW sleep and quiet wakefulness, claustral photo-stimulation induced a long-lasting silencing (up to 100-150 ms) of spike activity in almost all recorded FC units across layers in synchrony with large positive SWs (1-4 Hz) in the deep-layer LFP (Fig. 5c,d). The global silencing of units and associated LFP profile resembled the naturally-occurring Down-state of SW activity (Fig. 5c,d). Intriguingly, even a 5-ms photo-stimulation of the claustrum was sufficient to induce Down-state-like synchronous cortical silencing that lasted for over 100 ms. The claustrum-induced long-lasting silencing of cortical spike activity was behavioral state-dependent. That is, it was present only during quiet wakefulness and SW sleep, but absent during active wakefulness with phasic EMG activity (Fig. 5c,d). Claustral stimulation during active waking only resulted in brief (shorter than 50 ms) and partial suppression of cortical spike activity (Fig. 5c,d). To clarify the difference in claustral effects on neuronal subtypes, we classified cortical single units (121 units from 4 mice) into narrow-spike waveform (NS) units, which represent a subset of inhibitory interneurons, and wide-spike waveform (WS) units. Claustral stimulation drove spike response preferentially in NS units (8 / 16 units [50%]) compared with WS neurons (5 / 105 units [5%]), consistent with the results of the slice experiments (Fig. 3, Supplementary Fig. 4 and 5). Among the claustrum-driven units, NS units showed shorter spike latency than WS units, suggesting putative monosynaptic excitation of NS neurons following claustral activation (Supplementary Fig. 5). Immediately after the spiking of NS units, almost all units including their own were markedly suppressed. Following the long-lasting suppression, these units were reactivated synchronously (Fig. 5g,h). Firing rates immediately after the suppression were higher than those just before the suppression in a majority of units (Fig. 5i), suggesting that clausrum-induced SW regulation facilitates the synchronized firing of cortical neurons in the Down-to-Up state phase transition.

**Fig. 5.**
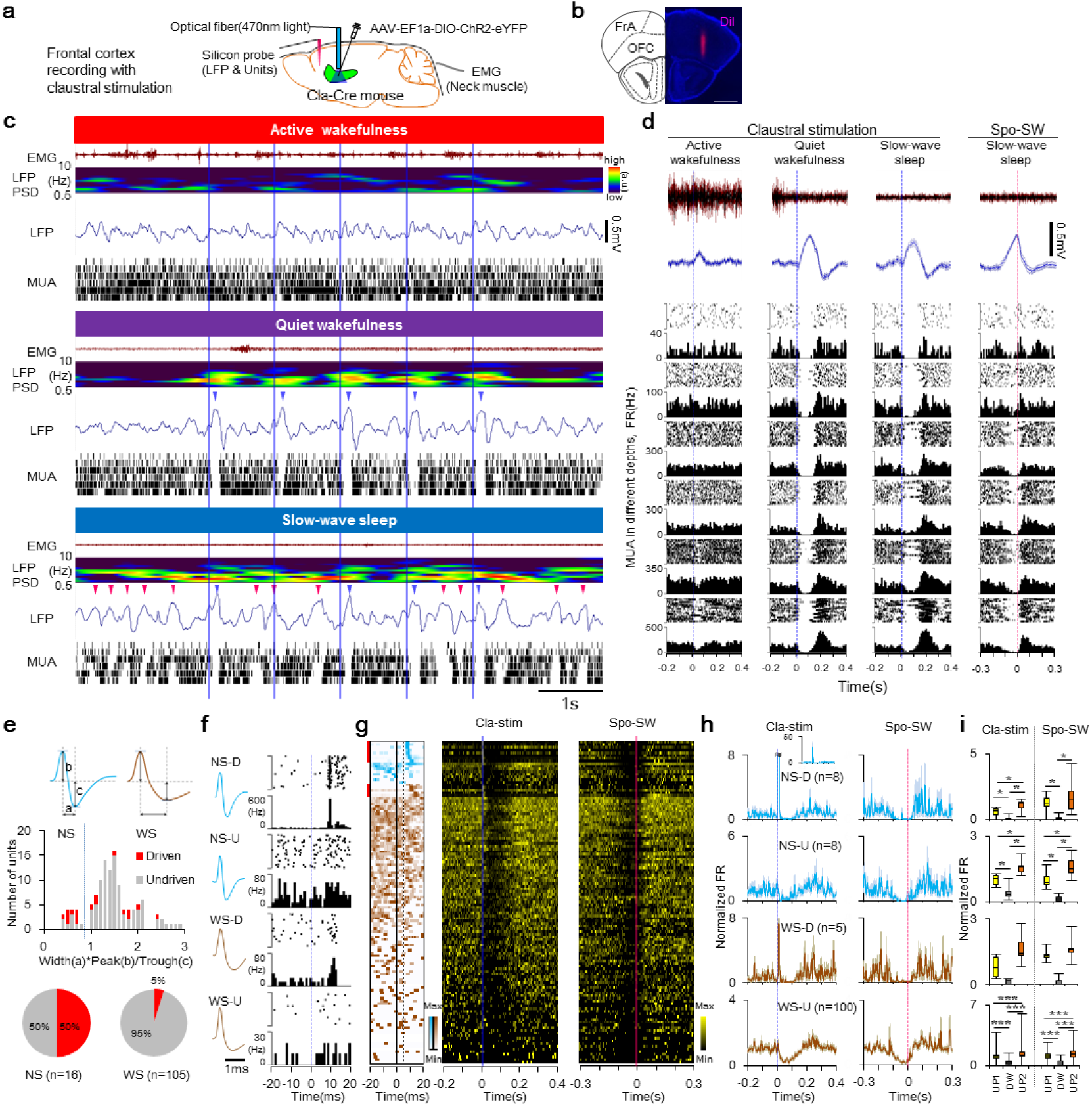
Optogenetic Stimulation of Claustrum Induces a Neocortical Down-State. **a**, Schematic of cortical recording with optogenetic claustral stimulation. **b**, An example recording track in frontal cortex (corresponding to data of **c, d**). Scale bar, 1 mm. **c**, Representative traces of LFP and multi-unit activities (MUAs) recorded from the frontal association and orbitofrontal cortex during active wakefulness (top), quiet wakefulness (middle) and SW sleep (bottom). Above these traces are shown the EMG and PSD of the LFP. Photo-stimulation (blue vertical line; 5 ms duration) was applied to the claustrum 5 times at 1 Hz. Each MUA was recorded from different cortical depths (approximately every 200 μm). Blue wedges indicate positive peaks of the claustrum-induced SWs. Magenta wedges indicate that of spontaneously occurred SWs. **d**, Photo-stimulation-triggered averages of EMG and LFP and PETHs of each MUA (20 traces, 10 ms bins) during active wakefulness (1st column), quiet wakefulness (2nd column), and SW sleep (3rd column). Averages of EMG and LFP and PETHs of MUA in the 4th column were calculated in reference to the positive peak of spontaneous SW. **e**, Classification of neocortical units into narrow-spike waveform (NS) units and wide-spike waveform (WS) units, and their responses to photo-stimulation of the claustrum. For each single unit, an index of waveform (width of peak to trough (ms) x ratio of peak/trough height) was calculated. Each unit was separated into either the NS or WS group by the trough of the histogram (the index value of 0.9). Units marked by red show spike responses to claustral stimulation, whereas units marked by gray did not. **f**, Examples of photo-stimulation-triggered PSTHs (1-ms bins) of each type of unit. Claustral stimulation-driven NS (NS-D), -undriven NS (NS-U), -driven WS (WS-U) and -undriven WS (WS-U) unit. **g**, Left column, PSTHs (1-ms bins, moving average of 3 bins) of single-unit spike activity during the period from 20 ms before and to 20 ms after the photo-stimulation. Time 0 indicates the onset of photo-stimulation. NS and WS units are shown by blue and brown, respectively. Claustral stimulation-driven units are marked with red bars. Middle column, PSTHs (1-ms bins, moving average of 7 bins) during the period from 0.2 s before to 0.4 s after the photo-stimulation. Right column, PETHs in reference to the positive peak of spontaneous SW (during the period from 0.3 s before to 0.3 s after the positive peak, 1-ms bins, moving average of 7 bins). Time 0 indicates the positive peak of spontaneous SW. Same units are aligned in the same rows across the left, middle and right columns. **h**, Left column, averaged spike activities of NS-D, NS-U, WS-D and WS-U units. The FR of each unit was normalized by its mean FR before averaging among the same type. **i**, Comparison of spike activity in the Up-state before the Down-state (UP1, yellow), the Down-state (DW, gray), the Up-state after the Down-state (UP2, red) of each unit type in **g**. Each FR was normalized as in **h**. For claustral stimulation, calculation periods of UP1, DW and UP2 were −0.18 to −0.08 s, 0.03 to 0.13 s and 0.18 to 0.28 s, respectively. Similary, for spontaneous SW, −0.25 to −0.15 s, −0.1 to 0 s and 0.05 to 0.15 s, respectively. Box plots indicates medians, quartiles (boxes) and ranges (minimum, maximum; whiskers). *P < 0.05, ***P < 0.001 (Two-sided Wilcoxon signed rank test with Holm’s correction).

Finally, we examined the cortical distribution of claustrum-induced SW activity. Simultaneous recordings of surface EEG from multiple cortical regions and LFP from prefrontal cortex (PFC) were performed during ChR2-mediated claustral stimulation (Fig. 6a). We found that claustral stimulation induced SW activity most strongly in the frontal cortex, moderately in the parietal cortex, and weakly in the occipital cortex (Fig. 6b-d). The cortical distribution of claustrum-induced SW activity varied among stimulation trials: in some cases it was localized within the frontal cortex, but in other cases, occured in almost the entire neocortex synchronously (Fig. 6b). This cortical distribution of claustrum-induced SW activity resembled that of the PFC-LFP positive peak-triggered average of spontaneous SW activity (Fig. 6b-d). These results suggest that the claustrum can regulate synchronized SW generation in widespread cortical areas through the synchronized and selective activation of cortical inhibitory interneurons.

**Fig. 6.**
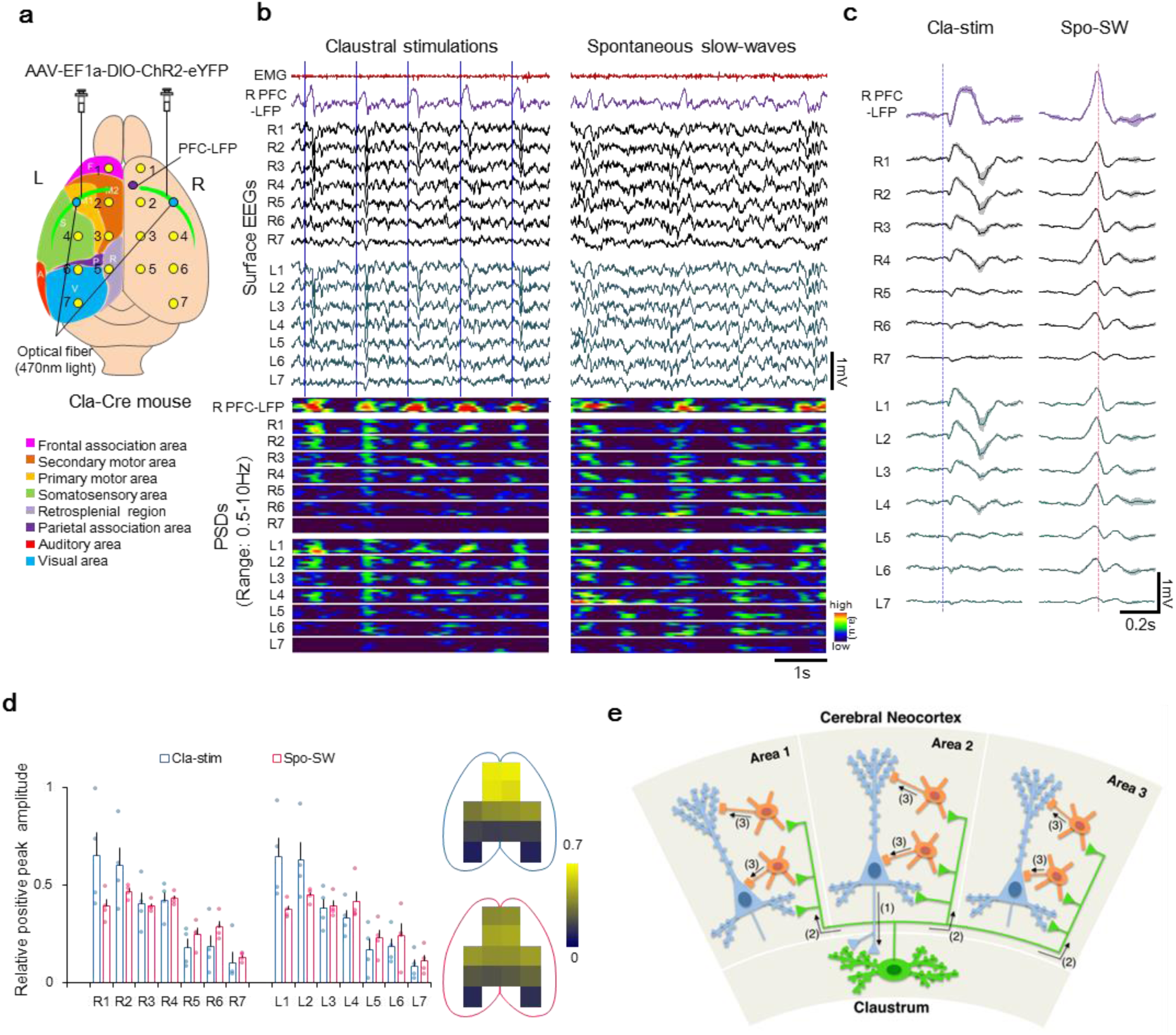
Optogenetic Stimulation of Claustrum Induces Widespread Slow-Wave Activity in Neocortex. **a**, Schematic of multiple EEG recordings from the neocortex and photo-stimulation of the claustrum. LFP in the deep layer of right prefrontal cortex (PFC) was also recorded to monitor local SWs. **b**, EEG recorded at each site of the cortical surface (R1-R7, L1-L7) during bilateral claustral stimulations (5-ms duration, 1 Hz, left column) and during the period that shows spontaneous SWs (right column) in the same recording session. Top two traces are EMG and LFP in the deep layer of the PFC. Heat maps are PSDs of the PFC-LFP and each EEG. **c**, Left column: Grand mean of PFC-LFP and EEG at each cortical site (R1-R7, L1-L7) in reference to the claustral photo-stimulation (n = 4 mice, 30 traces each). Right column: Grand mean of PFC-LFP and EEGs in reference to the positive peak of spontaneous SW in the PFC-LFP (n = 4 mice, 100 traces each). Error bars are s.e.m. **d**, Relative positive peak amplitude of the claustrum-induced SW (blue) and spontaneous SW (magenta) in averaged EEG at each site in **c**. Positive peak amplitudes of EEG SW were normalized within a mouse as that of PFC-LFP was 1. Although two-way repeated-measures ANOVA detected interaction effect (F(13,39) = 8.92, P < 0.001), *post hoc* test did not detect significant difference between groups at any recording site. Upper map with blue outline shows the spatial distribution of the claustrum-induced SW, while the lower magenda map indicates that of the spontaneous SW. Error bars are s.e.m. **e**, Schematic diagram of the neural circuit between the claustrum and neocortex. (1) Cre-expressing claustral neurons (green) receive excitatory inputs from pyramidal neurons (blue) in a given area of the neocortex (e.g. Area 2 in this figure). (2) The claustral neurons reciprocally and dispersedly send axons to almost all the excitatory and inhibitory neurons in widespread areas of the neocortex (e.g. Area 1, 2 and 3 in this figure) to induce EPSPs, but evoke action potentials predominantly in inhibitory interneurons (orange) in synchrony. (3) The synchronous firing of these inhibitory interneurons results in a prolonged silencing of most neocortical neurons, induction of a Down-state, and orchestration of neocortical SW activity.

## Discussion

Here, we describe for the first time the brain-wide functional connectivity and physiological effects of a major population of glutamatergic neurons in the clastrum. Cre-expressing claustral neurons make reciprocal connections with widespread cortical areas, especially in a large frontal network-containing higher-order association areas (*19*) such as the insular, prefrontal, cingulate, motor, and entorhinal cortices (c.f. also *6*). The cortical neurons projecting axons to the Cre-expressing claustral neurons are non-GABAergic and most of them display pyramidal cell morphology, indicating that the excitatory claustral neurons we identified receive glutamatergic excitatory inputs from the cortex. Within the claustrum, Cre-positive neurons receive massive inputs from both excitatory and inhibitory neurons that may enable intra-claustral signal propagation and information processing for subsequent output from the claustrum to the cortex (*13*). The most notable characteristic of the claustral output pattern is that the axons project dispersedly to a widespread network of higher-order association cortices and terminate in virtually all layers in each area, indicating dense coverage of cortical areas involved in cognitive processing. These observations suggest that the claustral neurons we studied do not transmit signals to specific target areas nor to selective target neurons, but rather send them non-selectively to essentially all higher-order areas and to all types of neurons (Fig. 6e). Indeed, slice electrophysiological experiments revealed that neurons in the insular and prefrontal cortices showed excitatory post-synaptic responses upon photo-stimulation of ChR2-expressing claustral neurons.

Although the activation of Cre-expressing claustral neurons elicited EPSPs in both excitatory pyramidal neurons and inhibitory interneurons in the cortex, their optogenetic activation drove suprathreshold action potentials predominantly in inhibitory interneurons, resulting in robust feed-forward inhibition of cortical activity that lasted for over 100 ms (Down-state). The present results demonstrate that the optogenetic-induced Down-state generation occured synchronously across multiple cortical areas and was followed by a synchronous Up-state which allows ensembles of neurons in multiple cortical areas to fire synchronously. The synchronous Up-state in multiple cortical areas during sleep is known to be critical for the coordinated replay of memory traces, successful transfer of memory-based information between cortical areas, and consolidation of long-term memory (*1-3*). Therefore, the population of excitatory claustral neurons may play a key role in coordinating the timing of a Down-state to multiple cortical areas to enable the following Down-to-Up transition to occur synchronously for inter-areal communication in SW sleep.

Crick and Koch proposed the hypothesis that the claustrum may represent a neural correlate of consciousness by orchestrating brain-wide cortical activity (*7*). In accord, it was reported that electrical stimulation of the claustrum results in a long-lasting inhibition of cortical neurons in the cat (*20, 21*) and reversible disruption of consciousness in an epileptic human patient (*22*). Our findings demonstrate that glutamatergic neurons in the claustrum regulate global neocortical SW activity by regulating interneuron firing in widespread cortical areas. The resultant Down-to-Up state transition may be necessary for offline cognitive processing that may contribute either directly or indirectly to consciousness states.

## Acknowledgements

We thank S. Tonegawa for valuable comments; C. Yokoyama for critical reading of the manuscript; I. R. Wickersham, H. S. Seung and E. M. Callaway for modified rabies virus, K. Deisserroth for AAV-EF1a-DIO-hCHR2(H134R)-EYFP: RIKEN BSI Research Resources Center for technical assistance; and members of Yoshihara lab for discussion. This work was supported by research funds from RIKEN to Y.Y. and grants-in-aid for Scientific Research on Innovative Areas “Memory Dynamism” (25115005) from the Ministry of Education, Culture, Sports, Science and Technology of Japan to Y.Y.

## Author Contributions

K.N., R.M., A.A., and Y.Y. conceived the study. S.M. and Y.Y. developed the Cla-Cre mouse. R.M., M.S., S.M, H.H. J.P.J., and Y.Y. performed anatomical experiments. A.A. performed *in vitro* electrophysiological experiments. K.N. performed *in vivo* electrophysiological experiments. K.N., R.M., A.A., K.M., and Y.Y. wrote the paper.

## Competing Interests

The authors declare no competing interests.

## Methods

### Animals

C57BL/6J background mice (ranging from 1 to 8 months old) were used for all the experiments. Mice were housed under standard housing conditions with food and water *ad libitum*. ACTB-tTA2-LacZ transgenic mice (strain Tg(ACTB-tTA2,-MAPT/lacZ)1Luo/J; stock number 014092) (*23*) were obtained from Jackson Laboratory. All animal experiments were approved by the Animal Care and Use Committee of RIKEN and conformed to the National Institutes of Health (NIH) guidelines.

### Generation of Cla-Cre Transgenic Mice

The Cla-Cre transgenic mouse line was established in the course of our olfactory circuitry research analyzing the enhancer/promoter region of mouse *Tbx21* gene which is specifically expressed in the mitral/tufted cells of the olfactory bulb. The 5.0-kb upstream sequence of mouse *Tbx21* gene (*12*) was fused to bacteriophage P1 Cre recombinase gene with human β-globin gene intron and Simian virus 40 (SV40) polyadenylation signal to generate *pTbx5.0-Cre* plasmid. *Tbx5.0-Cre* transgene was excised, gel-purified, and injected into the pronucleus of fertilized eggs that were obtained from crossing C57BL/6J and DBA/2J mice. The manipulated eggs were cultured to the two-cell stage and transferred into oviducts of pseudo-pregnant foster females (ICR strain). Genomic integration of the transgene was screened by PCR of tail DNA and transgene expression was examined by immunohistochemistry with anti-Cre antibody. Seven transgenic mouse lines were established and one of them (line #1) showed ectopic expression of Cre protein specifically in the claustrum (Fig. 1). This transgenic line was designated as the Cla-Cre mice, backcrossed with C57BL/6J mice more than ten times, and used in this study.

### Immunohistochemistry

Immunohistochemistry was performed as described previously (*24*). The following primary antibodies were used: mouse anti-Cre (1:1000, Millipore MAB3120), rat anti-GFP (1:1000, Nacalai Tesque GF090R), rabbit anti-DsRed (1:500, Clontech 632496), rabbit anti-GABA (1:1000, Sigma A2052), goat anti-Latexin (1:200, Acris TA303204), mouse anti-NeuN (1:500, biotin-conjugated, Millopore MAB377B), goat anti-choline acetyltransferase (1:400, Millipore AB144P), rabbit anti-serotonin (1/5000, Sigma S5545-2ML). Cy3- and Cy5-conjugated secondary antibodies were purchased from Jackson ImmunoResearch. Alexa488-conjugated secondary antibodies were purchased from Invitrogen. All the secondary antibodies were used at 1:400. Images were acquired with MVX10 fluorescence microscope (Olympus), Axio Imager Z1 (Zeiss), BZ-X700 (Keyence) or Fluoview FV1000 confocal laser-scanning microscope (Olympus).

### Viral Preparation

pAAV-TRE-sypGFP-2a-WGA plasmid was constructed by inserting synaptophysin-EGFP cDNA (*25*) and WGA cDNA (*26*) into pAAV-TRE-HGT plasmid (Addgene #27437). AAV5-TRE-sypGFP-2a-WGA (3.6 x10^12^ gp/ml) was custom-produced by University of North Carolina (UNC) Vector Core. The viral titer was estimated based on the quantitative PCR and indicated as genome particles (gp) per ml.

The following AAV vectors were purchased from UNC Vector Core:

- AAV5-CAG-FLEx-tdTomato (8.0 x 10^12^ gp/ml)
- AAV5-EF1a-FLEx-TVA-mCherry (3.0 x 10^12^ gp/ml)
- AAV8-CA-FLEx-RG (1.2 x 10^12^ gp/ml) The following AAV vector was purchased from University of Pennsylvania Vector Core.
- AAV5-EF1a-DIO-hChR2(H134R)-eYFP (1.9 – 2.6 x 10^12^ gp/ml)

Materials for generating modified rabies virus (RVΔG) were provided by Drs. Ian R. Wickersham, H. Sebastian Seung (MIT, MA) and Edward M. Callaway (Salk Institute, CA). EnvA-pseudotyped virus was produced as previously described (*27, 28*). The titer of the RVΔG was estimated to be 6.3 x 10^9^ IU/ml based on the infection efficiency of the 293T-TVA800 cell line.

### Viral Injection

To visualize the axonal projection of Cre-positive claustral neurons, 1 μl of AAV5-CAG-FLEx-tdTomato (8 x 10^12^ gp/ml) was injected into both anterior (AP: 1.78, ML: 1.5, DV: 2.45 mm) and posterior (AP: −0.46, ML: 3.3, DV: 2.63 mm) claustrum in the left hemisphere. The brains were analyzed 4 weeks after the viral injection.

For retrograde transsynaptic tracing using modified rabies virus, 1 μl of a 1:1 mixture of AAV5-CAG-FLEx-TVA-mCherry and AAV8-CA-FLEx-RG was injected into the left claustrum (AP: 0.5, ML: 3.1, DV: 2.63 mm). Three weeks later, 1 μl of RVΔG (pseudotyped with EnvA) was injected into the same brain location. The brains were analyzed 7 days after RVΔG injection. For quantification of axon terminals of Cre-positive neurons, Cla-Cre mice were crossed with ACTB-tTA2-LacZ mice to obtain Cre- and tTA-double-positive mice. These mice were injected at the left claustrum (AP: 0.5, ML: 3.1, DV: 2.63 mm) with 1.5 μl of AAV5-TRE-sypGFP-2a-WGA and analyzed after 6 weeks.

For slice electrophysiology, 0.75-1.0 μl of AAV5-EF1α-DIO-hChR2(H134R)-eYFP was injected into the left claustrum (AP: 0.5, ML: 3.1, DV: 2.63 mm) of Cla-Cre mice.

For in vivo electrophysiology, 1.5 μl of AAV5-EF1α-DIO-hChR2(H134R)-eYFP was injected into the right claustrum (claustral and cortical unit recording; AP: 0.5, ML: 3.1, DV: 2.6 to 2.7 mm) or bilateral claustrum (multiple EEG recording; AP: 1.0, ML: 3.0, DV: 2.5 mm) of Cla-Cre mice.

### Quantification of Input and Output Regions of Cre-positive Claustral Neurons

To quantify the axon terminals of Cre-positive neurons, every 6^th^ sections from the brains of AAV5-TRE-sypGFP-2a-WGA injected Cla-Cre/ACTB-tTA2-LacZ double-transgenic mice (n = 4) were immunostained with anti-GFP and anti-NeuN antibodies. Tiling images of entire sections were captured with confocal laser scanning microscope with 10x objective lens. Fluorescence intensities were analyzed using Image J software (National Institutes of Health). Boundaries of brain areas were registered manually according to Paxinos and Franklin’s the Mouse Brain in Stereotaxic Coordinates (*29*). sypGFP signals were adjusted so that the minimal and maximal intensities of the whole brain range between 0 and 255. Signals were then binarized (threshold = 210), and the fraction of pixels in each brain area was calculated. This value was normalized by the sum of pixels in total brain areas.

To quantify the presynaptic neurons of Cre-positive claustral neurons, every 6^th^ sections from the RVΔG injected mice (n = 4) were immunostained with anti-GFP and anti NeuN antibodies. The numbers of GFP-positive neurons in each brain area were counted manually. This number was normalized by the sum of total GFP-positive cells counted in the brain. All the quantifications were performed using data from the virus-injected hemisphere and did not contain the data of contralateral hemisphere.

### Slice Electrophysiology

ChR2-eYFP-expressing Cla-Cre mice (3-4 weeks after the virus injection) were deeply anesthetized with isoflurane, and transcardially perfused with an ice-cold N-methyl-D-glucamin (NMDG) solution composed of the following (in mM): 93 NMDG, 93 HCl, 30 NaHCO_3_, 25 glucose, 20 HEPES, 2.5 KCl, 1.2 NaH_2_PO_4_, 5 Na-ascorbate, 3 Na-pyruvate, 2 thiourea, 10 MgSO_4_, 0.5 CaCl_2_, 12 N-acetyl–L-cysteine, pH7.3, 315 mOsm. Following decapitation, horizontal or coronal slices (400 μm) were prepared on a vibratome (VT-1000s, Leica) in the same ice-cold NMDG solution, and incubated in warm (33° C) NMDG solution for 10 min. The slices were then transferred to warm (33° C) artificial CSF (ACSF) composed of the following (in mM): 124 NaCl, 2.5 KCl, 1.2 NaH_2_PO_4_, 24 NaHCO_3_, 5 HEPES, 12.5 glucose, 12 N-acetyl–L-cysteine, 2 MgSO_4_, 2 CaCl_2_, pH7.3, 315 mOsm (*30*) and then allowed to cool to room temperature. The same ACSF solution was also used for all subsequent recordings. All solutions were continuously bubbled with 95% O_2_ / 5% CO_2_. For recordings, slices were transferred to a submersion chamber on an upright microscope (Olympus BX51w1) and continuously superfused (3 ml/min) with oxygenated ACSF. Glass recording electrode (5-12 MΩ) was filled with a solution containing the following (in mM): 130 K-gluconate, 6 KCl, 4 NaCl, 10 HEPES, 2 EGTA, 4 MgATP, 0.3 Tris GTP, 10 phosphocreatine, 4% biocytin, pH 7.3, 300 mOsm. Whole-cell patch-clamp recordings were performed from various types of cortical neurons using a Multiclamp 700A patch amplifier (Molecular Devices) at room temperature in current-clamp mode controlled by custom-written routines in Igor Pro (WaveMetrics). All *in vitro* recordings were carried out from slices of the agranular insular or the prefrontal cortices.

For classification of the recorded neurons in cortices (fast-spiking and regular-spiking), firing patterns in response to current injection (10-500 pA) and measurement of single action potential’s half width were used (*16*). Additionally, following the electrophysiological experiments, slices were fixed in 4% PFA 0.15% picric acid in PBS. To visualize biocytin-filled cells, the slices were re-sectioned into 50-100 μm-thick slices after cryoprotection with 20% sucrose in PBS. The sections were blocked for 1 hour at room temperature in PBS containing 0.2% Triton X-100 and 10% normal horse serum, incubated in Alexa-Cy5-cojugated streptavidin (1:500; Invitrogen) for 16 - 20 hours, washed, mounted in Vectashield (Vector), and observed on a confocal microscope (FV1000, Olympus).

ChR2-expressing claustral neurons were photoactivated by blue LED light (∼470 nm; BRC or Doric Lense) through a optical fiber (Φ∼500 μm; BRC or Doric Lense). Brief light pulses (2-20 ms each, 8-17 mW) were delivered to the agranular insular or prefrontal cortices, while patch-clamp recordings were carried out from single cortical neurons.

### In vivo Electrophysiology

#### Surgery

For in vivo recording, adult male Cla-Cre mice (2 to 6 months old) were used. Surgery for electrodes implantations and viral injections was conducted under anesthesia with isoflurane (1-2%) or intraperitoneal injection of the mixture of medetomidine (0.3 mg/kg), midazolam (4.0 mg/kg) and butorphanol (5.0 mg/kg) (*31*).

For unit recording experiments, an electrical grand and a reference electrode were implanted above the cerebellum. An electroencephalogram (EEG) pin electrode was implanted above the parietal cortex (AP: −2.0, ML: 2.0 mm) contralateral to the unit recording side. Electromyogram (EMG) electrodes were inserted into neck muscle. For unit recordings with silicon probe, small cranial holes (Φ ∼2 mm) were made over the target areas: for claustrum (AP: 0.0 to 2.0, ML: 1.0 to 3.5 mm), for frontal cortex (AP: 1.5 to 3.0, ML: 1.0 to 2.0 mm). These holes were sealed with a silicone impression material (Shofu). For photo-stimulation of claustrum, an optical fiber cannula (Φ 400 μm) was implanted at a position just medial to the claustrum (AP: 0.5, ML: 2.8, DV: 2.6 mm from dura). The optical fiber cannula was inserted at a 30 degrees medial to lateral in most animals to enable easy access of recording probe into the claustrum. For head fixation, a head plate was affixed to the skull with adhesive.

For surface EEG recording from multiple cortical sites, totally 14 holes (Φ ∼1 mm) were made on the dorsal surface of the skull and pin electrodes (Φ 0.53 mm, impedance: 3-10 kΩ at 1 kHz) were implanted epidurally (Fig. 6a). Optical fiber cannulas (Φ 200 μm) were implanted bilaterally at the position dorsal to the claustrum (AP: 1.0, ML: 3.0, DV: 2.3 mm from dura). An additional wire electrode for LFP was implanted in medial PFC (AP: 2.0, ML: 1.0, DV: 1.0 mm).

After surgery, mice were singly-housed and allowed to recovery for 1 week before experiments.

#### Recordings

All recordings were performed during light phase of 12-h dark /12-h light cycle. In claustral and frontal cortex unit recordings, LFP and units activity were recorded using acutely inserted 32-ch silicon multi-channel microelectorode arrays with 50-µm spacing linear contact sites (impedance: 0.5-3 MΩ at 1 kHz) (Neuronexus). Electrical signals from the silicon probe and other electrodes were filtered (0.1-6000 Hz for LFP and EEG, 100-6000Hz for EMG) and sampled (at 15 kHz for LFP, EEG and EMG; at 30 kHz for spike activity) using 48-ch Cheetah recording system (Neuralynx).

On the recording, the head of mouse was fixed to the stereotaxic apparatus (Narishige) using the previously affixed head plate. Trunk of the animal was mildly restrained by covering with a plastic cylinder. The silicone material on the cranial windows was gently removed and locally anesthetized with drops of 1% lidocaine solution. The silicon probe was coated with DiI solution (Life technologies) for marking recording tracks and was slowly inserted into the target regions; claustrum (AP: 0.0 to 2.0 mm, ML: 1.0 to 3.5 mm, DV: 2.8 to 3.3 mm from dura), frontal cortex (AP: 1.5 to 3.0 mm, ML: 1.0 to 2.0 mm, DV: 1.5 to 3.0 mm from dura) according to the mouse brain atlas (Franklin & Paxinos, 2008). After inserting the silicon probe, the cranial window was filled with saline to avoid drying of the surface. To activate ChR2, we applied blue LED light (470 nm; Doric Lenses) through an optical fiber (Doric Lenses or Thorlab; Φ400 μm, 5-12mW at fiber tip) under the control of stimulator (Nihon Kohden). After a recording, cranial window was sealed with silicone impression material. The recording was repeated with the same animal with changing recording sites over a few weeks.

In EEG recording from multiple cortical sites, EEG and LFP signals were filtered at 0.1-6000 Hz and EMG at 100-6000 Hz using Cheetah recording system (Neuralynx) and all signals were sampled at 2 kHz by CED recording system (Cambridge Electronic Design). For photo-stimulation, blue LED light (470 nm; Doric Lenses) through optical fibers (Doric Lenses or Thorlab; Φ200 μm, 1.0-2.5 mW at fiber tip) was used.

#### Histology

After the recordings, animals were deeply anesthetized with pentobarbital and transcardially perfused first with phosphate-bufferd saline (PBS pH7.4) and then with 4% paraformaldehyde in PBS. The brains were removed, postfixed in 4% paraformaldehyde, immersed in 30% sucrose for cryoprotection, and then coronally sectioned into 50 or 100 μm thickness with a sliding microtome. Expression of ChR2-eYFP and recording tracks were observed with epifluorescence microscopy.

#### Data analysis

Spike2 software (Cambridge Electronic Design) was used for offline analyses of LFPs and unit activities including spike sorting and perievent time histograms. To reduce drifting noise of EEGs and LFPs, DC components of the signals were removed before analyses. Spikes were sorted mainly using principal component analysis.

#### Behavioral state definition

Based on EMG and EEG/LFP, we classified behavioral states into following 3 states.

- Active wakefulness: high and variable EMG and desynchronized EEG/LFP periods without spontaneous SW.
- Quiet wakefulness: transient low EMG periods (< 20 s) which occur between active wakefulness periods. Often lightly synchronized EEG/LFP (sporadic SWs) is accompanied.
- SW sleep: low EMG and synchronized EEG/LFP (continuous large SWs) periods which continues longer than 20 s.

#### Detection of SW-period

For detection of SW-periods in the claustral unit recording experiment (Fig. 4), EEG from parietal cortex was used. The EEG was downsampled to 200 Hz and band-pass-filtered at 1-20 Hz. The signal was transformed to power by calculating root mean square of the signal with 1-s time constant and smoothed. Using the power during an active wakefulness of 50 s in the same recording session as baseline SW power of non-SW-period, 1.3-1.5 times (adjusted depending on recording) of the baseline power was set as threshold for SW-period. Using this threshold as rising and falling threshold, SW-period-onset and -offset were automatically detected, respectively. SW-period-onsets which have at least 3 s of SW-period and SW-period-offsets which have at least 3 s of non-SW-period were used. Among these events, 23 to 30 events for each recording session were selected by visual inspection and used in PETH analysis in Fig. 4.

#### Detection of individual SW events

For detection of single SW events in the frontal cortex unit recordings (Fig. 5) and multiple EEG recordings (Fig. 6), LFP of the frontal cortex was used. The LFP was downsampled to 200 Hz and band-pass-filtered at 1-20 Hz. The standard deviation of the LFP during an active wakefulness of 50 s in the same recording session was calculated. Positive peaks which exceed 6 times of the SD were detected as positive peak of the single SW. For PETH analysis of frontal cortex single-units, 100 events of the SW were used.

#### Identification of claustral units

The identification of the claustral unit was carried out using opto-tagging by ChR2. Units recorded in the histologically verified recording track in the claustram were isolated into single units. To examine whether these units are directly driven by photo-stimulation (1 or 5 ms duration, 0.51 Hz), photo-stimulation-triggered PSTH (100 traces, 1-ms bins, 3 bins moving average) for each single unit was made (Fig. 4c). In this PSTH, baseline peak firing rate between 20 ms before and to the onset of photo-stimulation (−20 ms to 0 ms) was determined. If the peak firing rate within the 5 ms from the photo-stimulation onset (0 ms to 5 ms) exceeded twice of the baseline peak firing rate, the unit was classified as opto-tagged unit. In the case of units whose baseline peak firing rate was 0, at least 2 spikes within the 5 ms from the photo-stimulation was used as a threshold for opto-tagged unit. If peak firing rate within 5 ms from onset of photo-stimulation is equal to or lower than the baseline peak firing rate, these units were classified as untagged units. Among these untagged units, units recorded in the same or neighboring (50 µm apart) electrode to the opto-tagged units-detecting electrode was also analyzed as untagged neighboring claustral units. Other units that did not fit these criteria were not included in the analysis.

#### Criteria for claustrum-driven frontal cortex units

To examine whether frontal cortex units are driven by claustral stimulation, photo-stimulation-triggered PSTH (100 traces, 1-ms bins, 3 bins moving average) for each single unit was made (Fig. 5). In this PSTH, baseline peak firing rate between 20 ms before and to the onset of photo-stimulation (−20 ms to 0 ms) was determined. Because most of opto-tagged claustral units showed firing within 5 ms from the photo-stimulation onset (Fig. 4c), we set here time window of 5 ms to 15 ms for firing of cortical unit which is indirectly driven by photo-stimulation. If the peak firing rate during the 5 ms to 15 ms time window exceed twice of the baseline peak firing rate, the unit was classified as claustral-driven unit. In the case of units whose baseline peak firing rate is 0, at least 2 spikes within the 5 ms to 15 ms time window was used as a threshold for claustral-driven unit. Other units were classified as claustral-undriven unit.

#### Spike waveform classification

For the classification of neocortical neurons into narrow-spike waveform cells (NS, putative parvalbumin-positive inhibitory interneurons) and wide-spike waveform cells (WS, putative excitatory pyramidal cells and other types of inhibitory interneurons), an index of waveform (width of peak to trough (ms) x ratio of peak/trough height) was calculated. We defined here the low index value group (<0.9) as NS neurons and the high index value group (>0.9) as WS neurons (Fig. 5e).

## References and Notes

1. Ellenbogen, J. M., Payne, J. D. & Stickgold, R. The role of sleep in declarative memory consolidation: passive, permissive, active or none? Curr. Opin. Neurobiol. 16: 716–722 (2006).

2. Ji, D. & Wilson, M. A. Coordinated memory replay in the visual cortex and hippocampus during sleep. Nat, Neurosci. 10, 100–107 (2007).

3. Rasch, B. & Born, J. About sleep’s role in memory. Physiol. Rev. 93, 681–766 (2013).

4. Tononi, G. & Cirelli, C. Sleep and the Price of Plasticity: From Synaptic and Cellular Homeostasis to Memory Consolidation and Integration. Neuron 81, 12–34 (2014).

5. Steriade M. Neural substrates of sleep and epilepsy. (Cambridge Univ. Press, 2003)

6. Wang, Q. et al. Organization of the connections between claustrum and cortex in the mouse. J. Comp. Neurol. 525, 1317–1346 (2017).

7. Crick, F. C. & Koch, C. What is the function of the claustrum? Phil. Trans. R. Soc. B 360, 1271–1279 (2005).

8. Remedios, R., Logothetis, N. K. & Kayser, C. A role of the claustrum in auditory scene analysis by reflecting sensory change. Front. Syst. Neurosci. 8, 44 (2014).

9. Goll, Y., Atlan, G. & Citri, A. Attention: the claustrum. Trends in Neurosciences. 38, 486–495 (2015).

10. Kitanishi, T. & Matsuo, N. Organization of the claustrum-to-entorhinal cortical connection in mice. J. Neurosci. 37, 269–280 (2017).

11. White, M. G. et al. Anterior cingulate cortex input to the claustrum is required for top-down action control. Cell Rep. 22, 84–95 (2018).

12. Mitsui, S., Igarashi, K. M., Mori, K. & Yoshihara, Y. Genetic visualization of the secondary olfactory pathway in Tbx21 transgenic mice. Neural Syst. Circuits 1, 5 (2011).

13. Kim, J., Matney, C. J., Roth, R. H. & Brown, S. P. Synaptic organization of the neuronal circuits of the claustrum. J. Neurosci. 36, 773–784 (2016).

14. Arimatsu, Y., Kojima, M. & Ishida, M. Area- and lamina-specific organization of a neuronal subpopulation defined by expression of latexin in the rat cerebral cortex. Neuroscience 88, 93–105 (1999).

15. Callaway, E. M. & Luo, L. Monosynaptic circuit tracing with glycoprotein-deleted rabies viruses. J. Neurosci. 35, 8979–8985 (2015).

16. Connors, B. W. & Gutnick, M. J. Intrinsic firing patterns of diverse neocortical neurons. Trends Neurosci. 13, 99–104 (1990).

17. McCormick, D. A., Connors, B. W., Lighthall, J. W. & Prince, D. A. Comparative electrophysiology of pyramidal and sparsely spiny stellate neurons of the neocortex. J. Neurophysiol. 54, 782–806 (1985).

18. The Petilla Interneuron Nomenclature Group (PING) et al. Petilla terminology: nomenclature of features of GABAergic interneurons of the cerebral cortex. Nat. Rev. Neurosci. 9, 557–568 (2008).

19. Barbas, H. General cortical and special prefrontal connections: principles from structure to function. Annu. Rev. Neurosci. 38, 269–289 (2015).

20. Tsumoto, T. & Suda, K. Effects of stimulation of the dorsocaudal claustrum on activities of striate cortex neurons in the cat. Brain Res. 240, 345–349 (1982).

21. Salerno, M. T., Cortimiglia, R., Crescimanno, G., Amato, G. & Infantellina, F. Effects of claustrum stimulation on spontaneous bioelectrical activity of motor cortex neurons in the cat. Exp. Neurol. 86, 227–239 (1984).

22. Koubeissi, M. Z., Bartolomei, F., Beltagy, A. & Picard, F. Electrical stimulation of a small brain area reversibly disrupts consciousness. Epilepsy Behav. 37, 32–35 (2014).

## References in Methods

23. Miyamichi, K. et al. Cortical representations of olfactory input by trans-synaptic tracing. Nature 472, 191–196 (2011).

24. Yoshihara, S., Omichi, K., Yanazawa, M., Kitamura, K. & Yoshihara, Y. Arx homeobox gene is essential for development of mouse olfactory system. Development 132, 751–762 (2005).

25. Miyasaka, N. et al. From the olfactory bulb to higher brain centers: genetic visualization of secondary olfactory pathways in zebrafish. J. Neurosci. 29, 4756–4767 (2009).

26. Yoshihara, Y. et al. A genetic approach to visualization of multisynaptic neural pathways using plant lectin transgene. Neuron 22, 33–41 (1999).

27. Wickersham, I. R., Finke, S., Conzelmann, K. K. & Callaway, E. M. Retrograde neuronal tracing with a deletion-mutant rabies virus. Nat. Methods 4, 47–49 (2007).

28. Wickersham, I. R., Sullivan, H. A. & Seung, H. S. Production of glycoprotein-deleted rabies viruses for monosynaptic tracing and high-level gene expression in neurons. Nat. Protoc. 5, 595–606 (2010).

29. Franklin, K. B. J. & Paxinos G. The Mouse Brain in Stereotaxic Coordinate. Third edition, (Academic Press, 2008)

30. Ting, J. T., Daigle, T. L., Chen, Q., Feng, G. “Acute brain slice methods for adult and aging animals: application of targeted patch clamp analysis and optogenetics” in Patch – Clamp Methods and Protocols, Methods in Molecular Biology, Martina, M., Taverna, S. Eds. (Springer, New York, 2014), vol. 1183, chap. 14, pp. 221–242.

31. Kawai, S., Takagi, Y., Kaneko, S. & Kurosawa, T. Effect of three types of mixed anesthetic agents alternate to ketamine in mice. Exp. Anim. 60, 481–487 (2011).

