## Supplementary Materials for "The Claustrum Coordinates Cortical Slow-Wave Activity"

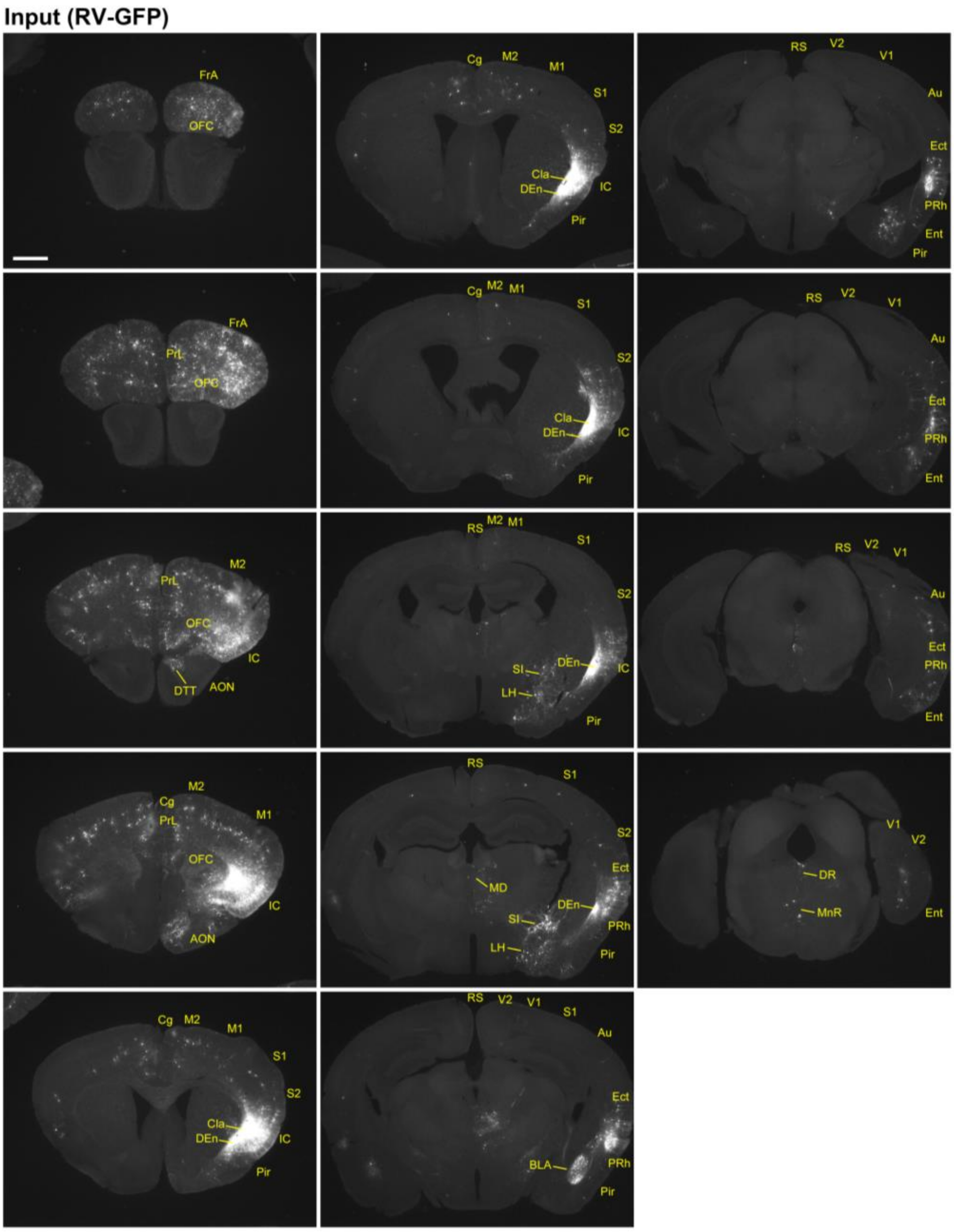


**Supplementary Figure 1**

**A Representative Example of Presynaptic Input Neurons to Cre-expressing Claustral Neurons.**

Presynaptic input neurons to Cre-expressing claustral neurons were visualized with a modified rabies virus-mediated mono-synaptic retrograde tracing method. AON, anterior olfactory nucleus; Au, auditory cortex; BLA, basolateral amygdala; Cg, cingulate cortex; Cla, claustrum; DEn, dorsal endopiriform nucleus; DR, dorsal raphe nucleus; DTT, dorsal tenia tecta; Ect, ectorhinal cortex; Ent, entorhinal cortex; FrA, frontal association cortex, IC, insular cortex; LH, lateral hypothalamic area; M1, primary motor cortex; M2, secondary motor cortex; MD, mediodorsal thalamic nucleus; MnR, median raphe nucleus; OFC, orbitofrontal cortex; Pir, piriform cortex; PrL, prelimbic cortex; PRh, perirhinal cortex; RS, retrosplenial cortex; S1, primary somatosensory cortex; S2, secondary somatosensory cortex; SI, substantia innominate; V1, primary visual cortex; V2, secondary visual cortex. Scale bar, 1 mm.

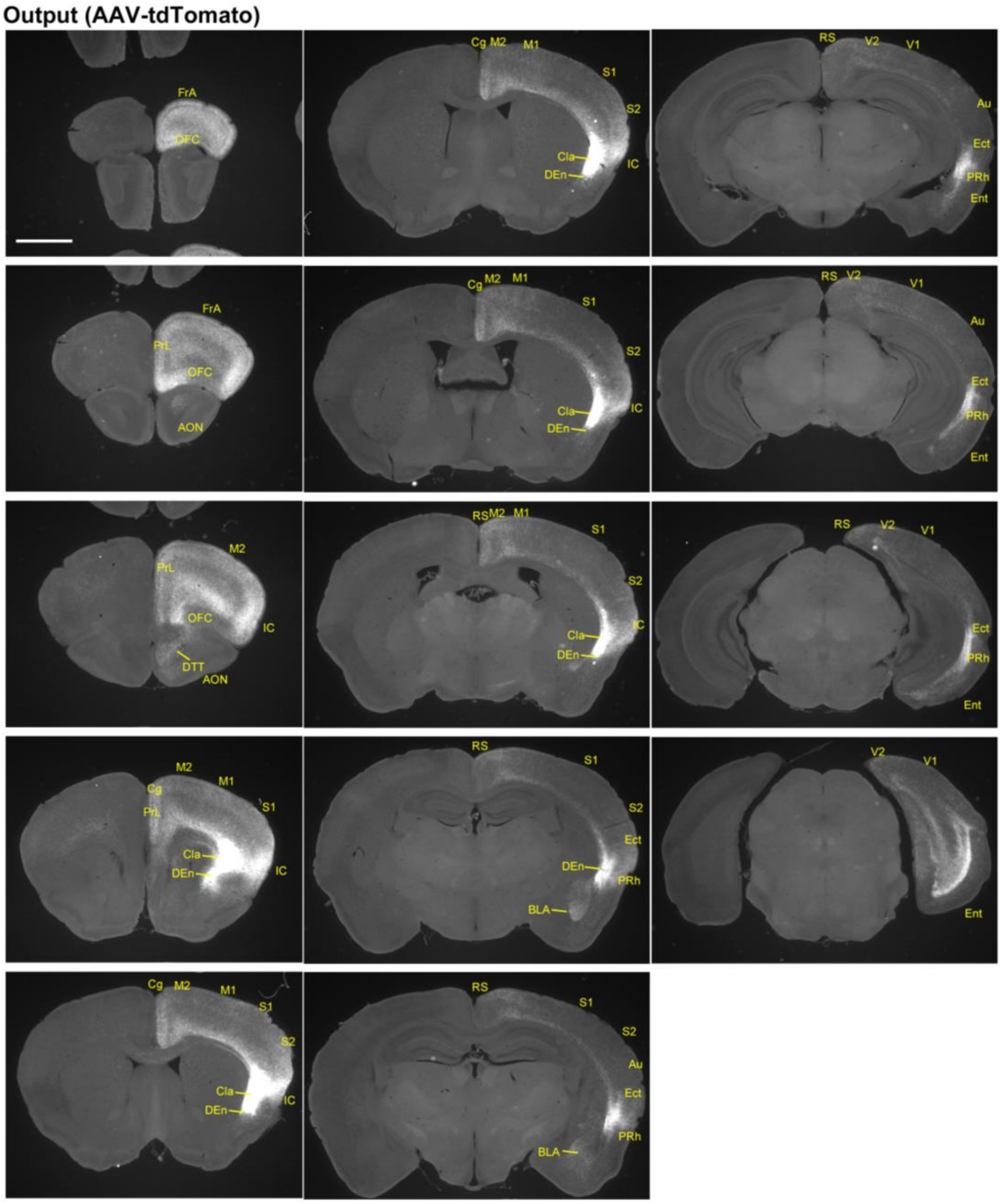


**Supplementary Figure 2**

**A Representative Example of Axonal Trajectories of Cre-expressing Claustral Neurons.**

Axonal trajectories of Cre-expressing claustral neurons were visualized with Cre-dependent tdTomato-expressing AAV. AON, anterior olfactory nucleus; Au, auditory cortex; BLA, basolateral amygdala; Cg, cingulate cortex; Cla, claustrum; DEn, dorsal endopiriform nucleus; DTT, dorsal tenia tecta; Ect, ectorhinal cortex; Ent, entorhinal cortex; FrA, frontal association cortex, IC, insular cortex; M1, primary motor cortex; M2, secondary motor cortex; OFC, orbitofrontal cortex; PrL, prelimbic cortex; PRh, perirhinal cortex; RS, retrosplenial cortex; S1, primary somatosensory cortex; S2, secondary somatosensory cortex; V1, primary visual cortex; V2, secondary visual cortex. Scale bar, 1 mm.

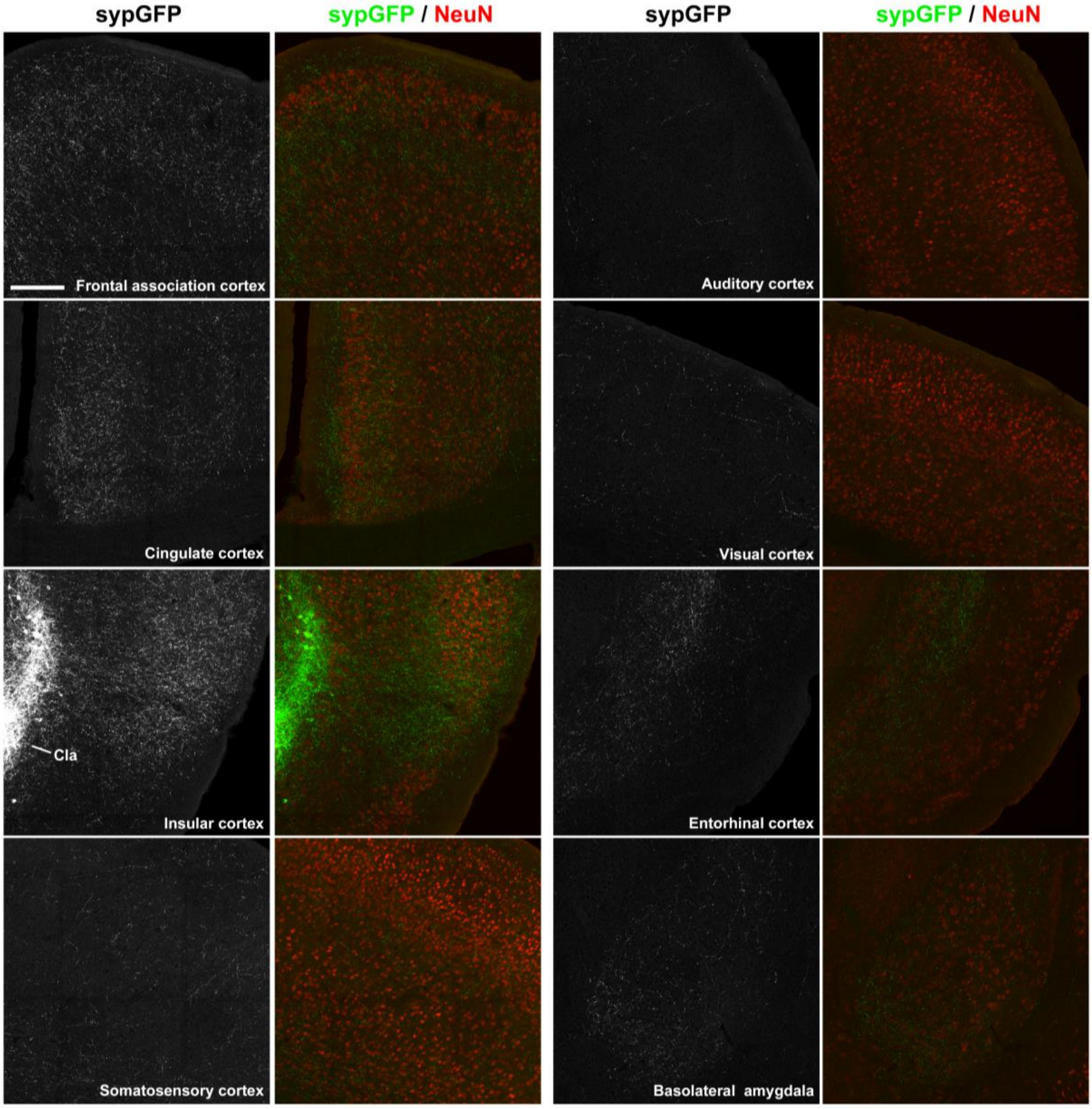


**Supplementary Figure 3**

**A Representative Example of Synaptic Vesicle Distributions of Cre-expressing Claustral Neurons.**

Synaptic vesicle distributions of Cre-expressing claustral neurons were visualized with Cre-dependent synaptophysin-GFP-expressing AAV. Scale bar, 200μm.

.

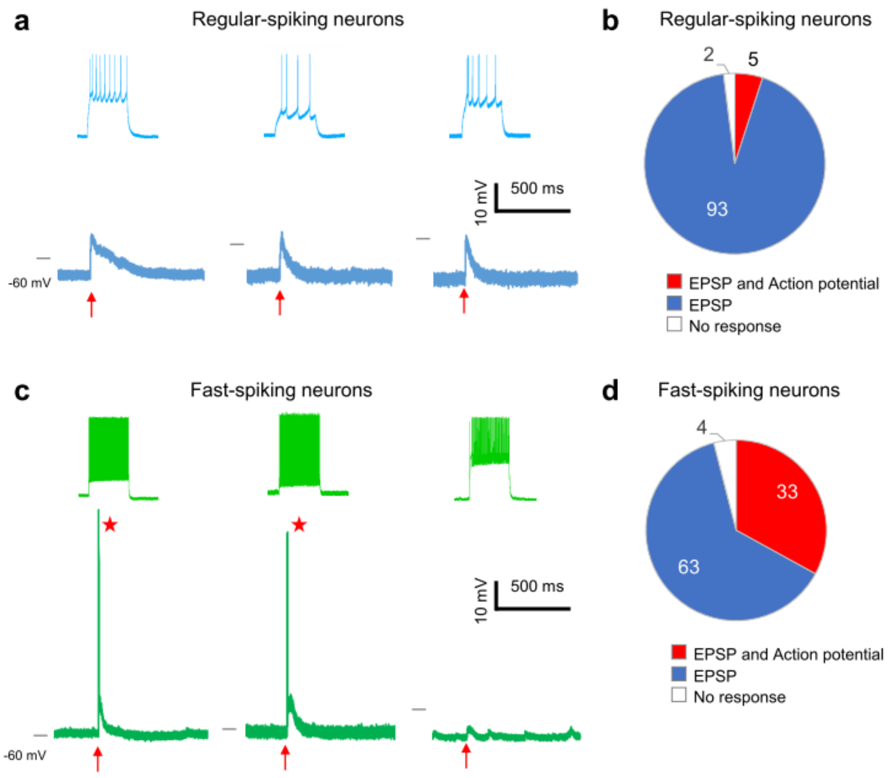


**Supplementary Figure 4**

**Claustral Stimulation Drives Fast-Spiking Interneurons in the Frontal cortex.**

**a**, Current-clamp recordings of regular-spiking neurons in frontal cortex. Firing patterns induced by inward current pulse injection (top), and responses to optogenetic stimulation of claustral axons (bottom).

**b**, Percentage of regular-spiking neurons (n = 90) that showed EPSP with action potential (red), EPSP without action potential (blue), and no response (white) upon optogenetic claustral stimulation.

**c**, Recordings from fast-spiking neurons.

**d**, Percentage of fast-spiking neurons (n = 27) that showed an EPSP with action potential (red), EPSP without action potential (blue), and no response (white) upon the optogenetic claustral stimulation.

Black bar on the left of each voltage trace, -60 mV; red arrow below each trace, light stimulation (5 ms); red asterisk, action potential.

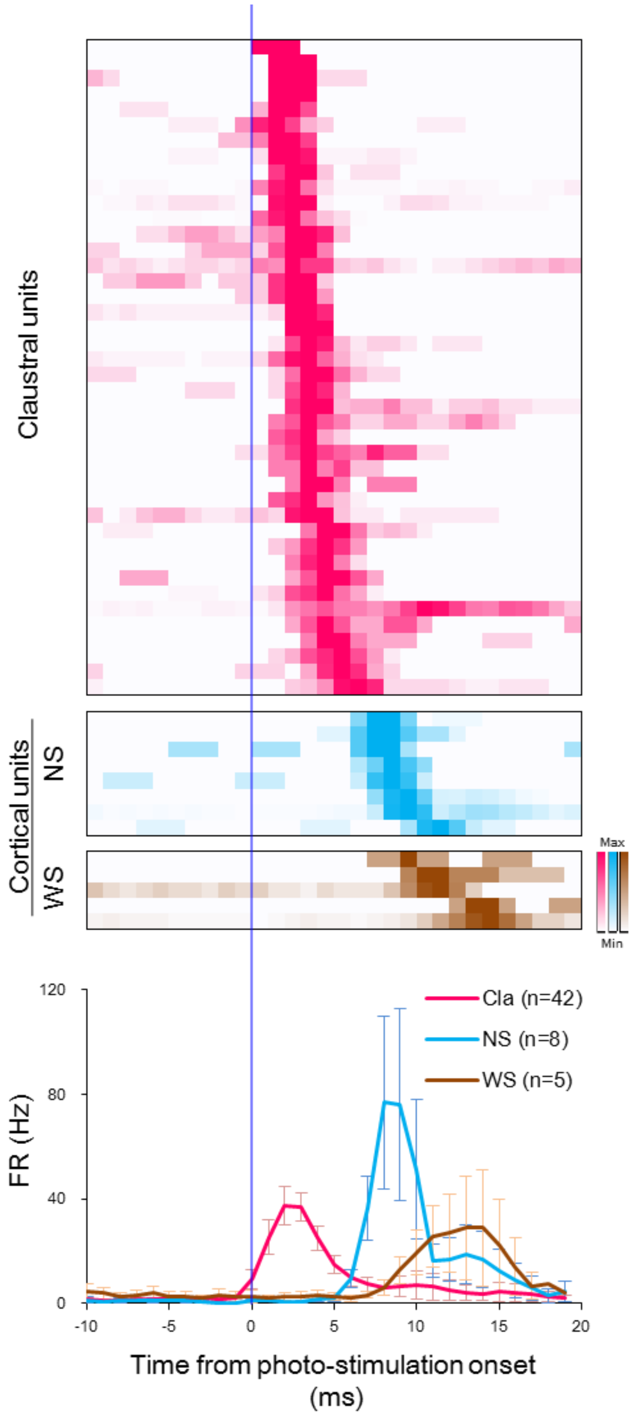


**Supplementary Figure 5**

**Spike Latencies of Claustral and Cortical Neurons after Photo-stimulation.**

Top, Claustral photo-stimulation-triggered PSTHs (1-ms bins, smoothed by 3 bins moving average) of claustral units (magenta), cortical narrow-spike wavefrom units (NS, blue) and cortical wide-spike waveform units (WS, brown). Bottom, group average of each PSTH. Error bars are s.e.m.
